# Current and future ocean chemistry negatively impacts calcification in predatory planktonic snails

**DOI:** 10.1101/2020.08.04.236166

**Authors:** Deborah Wall-Palmer, Lisette Mekkes, Paula Ramos-Silva, Linda K. Dämmer, Erica Goetze, Karel Bakker, Elza Duijm, Katja T.C.A. Peijnenburg

## Abstract

Planktonic gastropods mediate an important flux of carbonate from the surface to the deep ocean. However, we know little about the response of atlantid heteropods, the only predatory, aragonite shelled zooplankton, to ocean acidification (OA), and they are not incorporated in any carbonate flux models. Here we quantify the effects of OA on calcification and gene expression in atlantids across three pH scenarios: mid-1960’s, ambient, and future 2050 conditions. Atlantid calcification responses to decreasing pH were negative, but not uniform, across the three scenarios. Calcification was reduced from mid-1960s to ambient conditions, and longer shells were grown under 2050 conditions. Differential gene expression indicated a stress response at both ambient and future conditions, with down-regulation of growth and biomineralization genes with decreasing pH. Our results suggest that ocean chemistry in the South Atlantic is already limiting atlantid calcification, and that exposure to near-future OA triggers rapid shell growth under stress.

## Introduction

Calcifying planktonic gastropods play an important role in ocean carbonate flux, by transporting inorganic carbon from the ocean surface into deep waters via the rapid sinking of their relatively heavy calcium carbonate shells ^1–3^. Although small (up to∼2 cm), holoplanktonic gastropods are widespread and can be highly abundant in the upper ocean, exceeding densities of 10,000 individuals per m^3^ of seawater ^4^. Consequently, it is estimated that shelled pteropods, one holoplanktonic gastropod group, produce up to 89% of all pelagic calcium carbonate (CaCO_3_) ^3^, generating at least 12% of the total carbonate flux worldwide ^5^, and up to 33% of CaCO_3_ exported into shallow (∼100 m) waters ^3^. The transport of carbon from the ocean surface into deep water by planktonic gastropods (and other calcifying plankton) in turn allows the ocean to absorb more carbon, in the form of carbon dioxide (CO_2_). The oceans are therefore an important sink for CO_2_ and have taken up ∼30% of all cumulative releases of anthropogenic CO_2_, slowing the accumulation of CO_2_ in our atmosphere ^6^. However, the current increased uptake of CO_2_ by the oceans has led to a reduction in ocean pH at a rate unprecedented during the last 66 million years ^6–11^. The adverse consequences of this anthropogenic ocean acidification (OA) are being felt by many marine organisms ^12,13^. Recent research has confirmed negative effects of OA for shelled pteropods, including reduced calcification, increased shell dissolution and differential gene expression ^14–17^ and has highlighted them as useful OA-indicators, especially at higher latitudes ^15,18^. However, another abundant and ecologically important holoplanktonic gastropod group ^19^, the atlantid heteropods, have been largely overlooked in OA research. Apart from the physical structure of their shells ^20^, the calcification mechanisms for atlantids are unknown ^20,21^ and atlantids have not been considered in any models of carbonate flux thus far ^3^.

Pteropods and heteropods (holoplanktonic gastropods) are thought to be amongst the most susceptible groups to OA and likely the first to experience it, due to a combination of three factors that make these two groups similarly vulnerable. First, all holoplanktonic gastropods rely on an aragonitic shell; even species generally considered shell-less have a shell at the larval stage ^22^. Aragonite, a metastable form of calcium carbonate that is especially soluble in seawater ^23^, becomes difficult and energetically costly ^13,24^ to produce under OA conditions, and aragonitic shells can dissolve if aragonite undersaturation occurs ^25^. Second, holoplanktonic gastropods inhabit the upper ocean, where the greatest proportion of anthropogenic CO_2_ is being absorbed ^26^. Most holoplanktonic gastropods also undergo diel vertical migrations over hundreds of meters ^22,27^. With shoaling of the aragonite saturation horizon, they are increasingly likely to encounter deep waters that are undersaturated with respect to aragonite ^28^, thereby experiencing altered ocean chemistry across the vertical extent of their distributions. Third, holoplanktonic gastropods can have high abundances in cold, high latitude regions that have a higher capacity to absorb atmospheric CO_2_, thus more rapidly becoming acidic compared to warmer regions^28,29^.

Although superficially similar, shelled pteropods (Thecosomata) and atlantid heteropods (Atlantidae, Pterotracheoidea) have evolutionarily independent origins and occupy different trophic levels. While shelled pteropods are particle-feeders via mucous webs, juvenile atlantids feed on algae using a ciliated velum and adult atlantids are selective, visual predators. As such, atlantids are the only aragonite shelled predatory plankton, and are uniquely positioned to indicate the effects of changing ocean chemistry on higher trophic levels. Adult atlantids rely on shelled pteropods as a primary food source ^22^, which could make them even more vulnerable to the effects of OA. Atlantids are found in the epipelagic zone of open waters mainly from tropical to temperate latitudes, although there are two cold water species ^30^. *Atlanta ariejansseni* is the most southerly distributed atlantid species, being restricted to the Southern Subtropical Convergence Zone between 35–48°S, where it can reach abundances of up to ∼200 individuals per 1000 m^3^ and likely represents an important predator within the plankton ^31^. The cold water distribution and relatively high abundance of *A. ariejansseni* make it an excellent candidate as an OA sentinel species. In addition, juvenile atlantids are relatively easily maintained under laboratory conditions because they feed on algae using their ciliated velum. This makes them ideal organisms with which to study the effects of changing ocean carbonate chemistry on planktonic gastropod calcification.

Here we present results of the first growth and OA experiments focussed on heteropods, and the first transcriptome of a heteropod. We address the following questions: (1) What is the rate of atlantid shell growth under current ambient conditions in the South Atlantic Ocean? (2) Has atlantid calcification already altered in response to recent changes in high latitude ocean carbonate chemistry? (3) Will future ocean conditions lead to a decline in atlantid calcification, similar to the response found for shelled pteropods? Experiments were carried out in the South Atlantic Ocean on board the *RRS Discovery*. The calcification response of juvenile *A. ariejansseni* to variations in ocean carbonate chemistry was investigated under ambient conditions for up to 11 days (ambient shell growth), and under realistic past, ambient and future ocean carbonate chemistry scenarios for three days. We use a thorough multi-disciplinary approach, combining fluorescence microscopy (n=184) and micro-CT scanning (n=43) of the same individuals to quantify shell growth, as well as RNA sequencing of pooled individuals from the same experiments to detect differential gene expression as an OA response. Shell growth parameters show for the first time that ambient seawater conditions in the South Atlantic already limit atlantid calcification, and that predicted future ocean carbonate chemistry causes atlantids to grow faster, which may be a stress response. Shell growth measurements are supported by differentially expressed genes that indicate the down-regulation of growth and biomineralisation genes with increasing pH. This study demonstrates the suitability of atlantids as an OA sentinel, and highlights that changes in calcification and consequently a likely reduction in the transport of carbonate from the surface to the deep ocean, are already occurring in the South Atlantic Ocean.

## Results and Discussion

### Shell growth under ambient conditions

To measure shell calcification rate under current ocean conditions, juvenile *A. ariejansseni* collected from the Southern Subtropical Convergence Zone in the South Atlantic were stained with calcein indicator (Fig. 1a, 1e), incubated in ambient waters with high food availability and a subsample of specimens was collected every 2–3 days for up to 11 days. Here we show that the calcein indicator, which is incorporated into the shells during growth and can be detected using fluorescence microscopy, was only integrated into the apertural/growing edge of the shell, suggesting that the shell is not thickened from inside as is observed in some pteropods ^32^. Shell extension is therefore an informative measure of shell growth. Some small repairs from the inside surface of the shell, similar to those found in pteropods ^33^ were also observed (Supplementary Figure S1).

**Figure 1.**
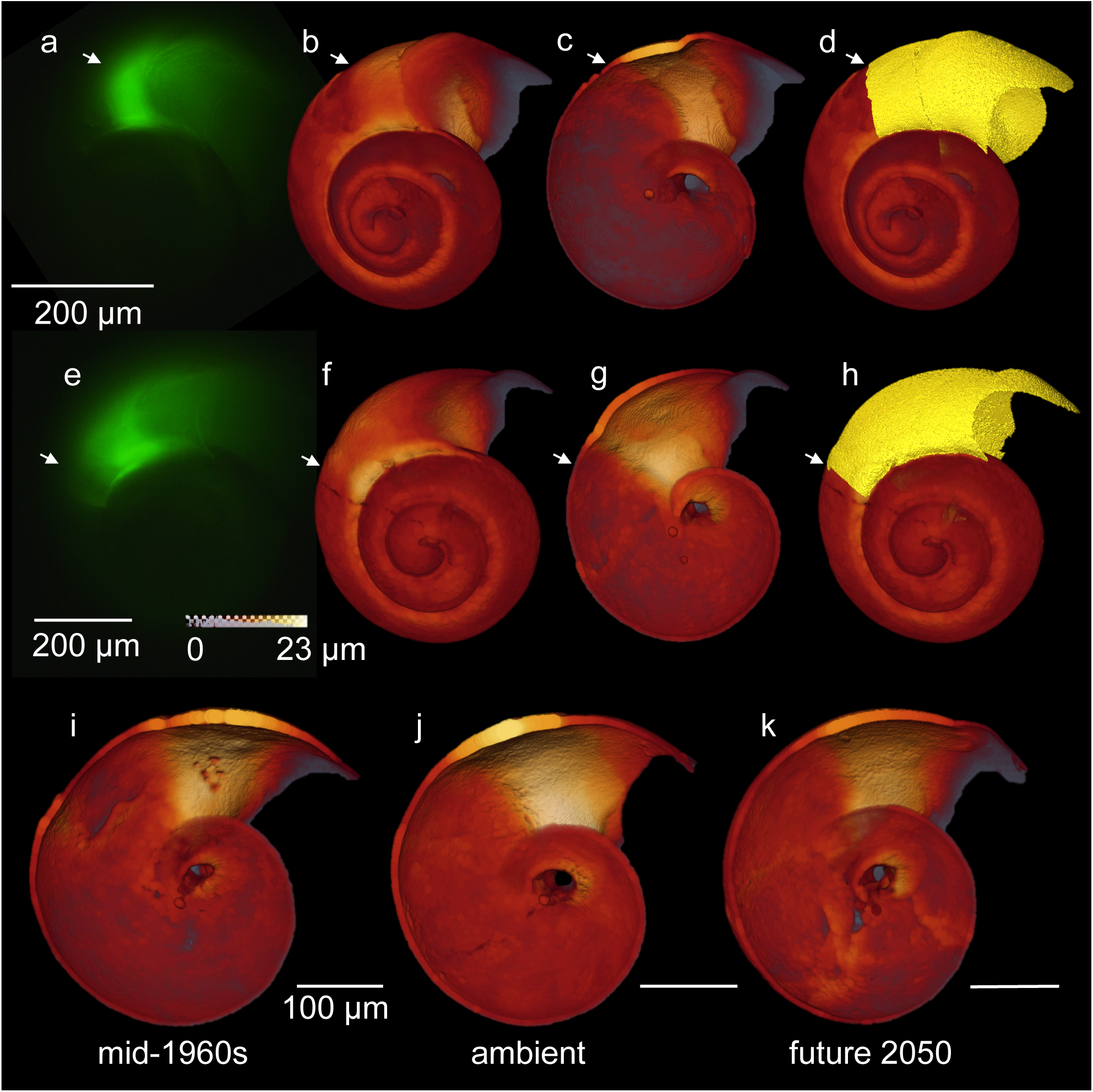
Quantifying calcification of *Atlanta ariejansseni*. (**a, e**) Shell extension of the shell was measured using the fluorescence images. (**b, c, f, g**) The position of the onset of the experiment/glow was then identified in thickness maps produced using micro-CT. White arrows show the onset of the experiment. Thickness maps presented in the ‘glow’ colour scheme. (**d, h**) The shell was segmented to isolate the part of the shell grown during the experiment. (**a-d, i**) mid-1960s, (**e-h, j**) ambient, (**k**) future 2050.

Over three to eleven days, specimens grew up to 99 µm (shell extension) per day (n=44, Table 1). Individuals grew significantly between sampling days (Kruskal-Wallis H(2)=12.08, p=0.017). However, there is a drop in mean shell extension at day nine, which may be due to a pause in shell growth to undergo metamorphosis (Table 1, Fig. 2a). Towards the end of the experiment, several individuals were observed to have undergone metamorphosis, having lost their velum and developed their swimming fin. Mean shell extension (all specimens) and maximum shell extension (largest specimen for each collection day) varied from 30–69 µm and 54–99 µm per day respectively, and both were found to decrease exponentially with age (Fig. 2b). This pattern may be due to the relatively broader surface of the shell as the atlantid increases with age, such that the amount of shell produced may be approximately the same. Adult specimens of *A. ariejansseni* exhibit the lowest growth rate of ∼25 µm per day (mean of 5 adult specimens). Assuming that the shell of *A. ariejansseni* follows the exponential decrease in shell extension identified by the mean shell extension per day (−26.89ln[days]+95.076), and assuming that the rate never falls below 25 µm per day, it would take around 116 days for an *A. ariejansseni* specimen to grow to full adult size (∼3200 µm of shell extension measured along whorl suture).

**Table 1.** Typical shell growth of *Atlanta ariejansseni* at ambient conditions over eleven days. N is the number of specimens sampled on each day.

| Sampling day | N | Measured DIC (μmol/kg) | Calculated TA (μmol/kg) | Measured pH (NBS) | Calculated pCO<sub>2</sub> | Calculated ΩAr | Mean total shell extension (μm) ± 1SD | Maximum total shell extension (μm) | Mean shell extension per day (μm) | Maximum shell extension per day (μm) |
| --- | --- | --- | --- | --- | --- | --- | --- | --- | --- | --- |
| 0 | - | 2112 | 2358 | 8.15 | 426 | 2.73 | - | - | - | - |
| 3 | 8 | 2130 | 2345 | 8.15 | 417 | 2.37 | 208 ± 65 | 299 | 69 | 100 |
| 5 | 10 | 2141 | 2368 | 8.17 | 399 | 2.49 | 231 ± 77 | 374 | 46 | 75 |
| 7 | 10 | 2135 | 2360 | 8.17 | 394 | 2.45 | 306 ± 87 | 456 | 44 | 65 |
| 9 | 9 | 2151 | 2364 | 8.14 | 426 | 2.35 | 271 ± 109 | 486 | 30 | 54 |
| 11 | 7 | 2173 | 2378 | 8.14 | 437 | 2.29 | 408 ± 148 | 599 | 37 | 54 |

**Figure 2.**
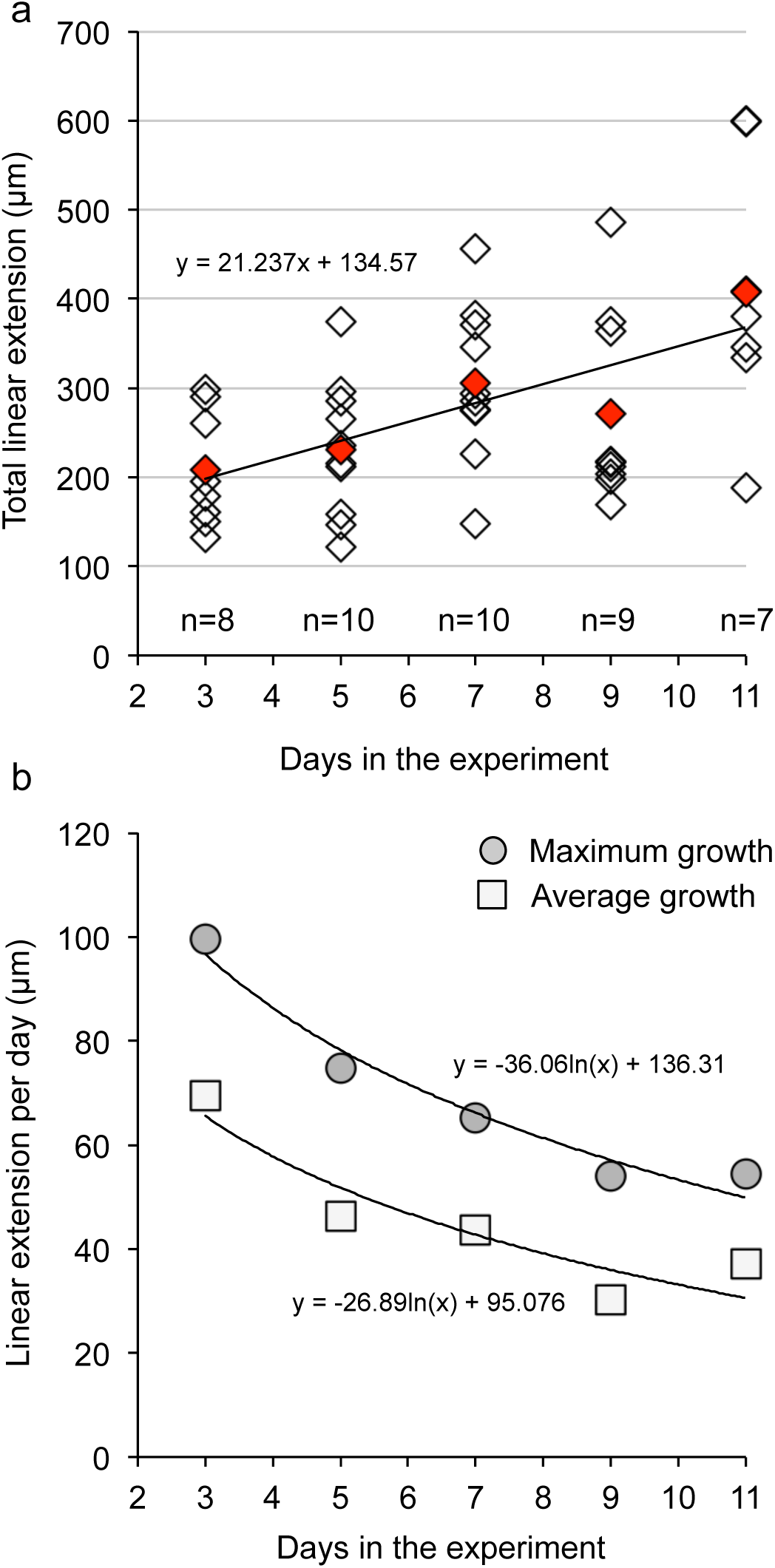
Shell extension of *Atlanta ariejansseni* in ambient conditions over 11 days. (**a**) Total shell extension of the shell. Red diamonds represent mean values for each sampling day. (**b**) Maximum and mean shell extension of the shell for each sampling day decreases exponentially and can be used to estimate the time taken to grown to full adult size, assuming constant conditions.

The tempo of the atlantid life cycle is, until now, completely unknown and our results give a first estimate of their minimum longevity. If atlantids reach reproductive maturity in ∼116 days, this could allow for more than one generation per year. This is comparable to the shelled pteropods, which are thought to live for ∼1–2 years and may produce two generations of offspring per year in the Southern Ocean ^34,35^. During specimen collection in the South Atlantic Ocean for the present study, small juvenile and large adult specimens were present in the same location at the same time, supporting this inference. Juvenile specimens of *A. ariejansseni* have been caught at the beginning (September) and end (February) of the summer growing season in sediment traps moored in the Southern Ocean offshore of Tasmania (47°S, 142°E) ^36^. This suggests that, similar to the pteropod *Limacina helicina antarctica, A. ariejansseni* could have an overwintering juvenile population.

### The effects of OA on calcification

To evaluate the effects of future and past ocean carbonate chemistry on atlantid calcification, specimens of *A. ariejansseni* were incubated for three days across three pH scenarios (n=184). We applied a past scenario of 0.05 pH units higher than ambient (ambient pH 8.14 ± 0.02, past pH 8.19 ± 0.02) that is approximately equivalent to the mid-1960s (assuming a decrease in pH of 0.001 units per year in this region) ^11^, and a future OA scenario of 0.11 pH units lower than ambient (pH 8.03 ± 0.00), which is approximately equivalent to expectations for the year 2050 in the South Atlantic Ocean (under IPCC Representative Concentration Pathway RCP8.5) ^7,37^. Aragonite saturation was maintained in all scenarios (Ω>1.82). Calcification was measured in three ways: mean shell extension was measured from fluorescence images, while the volume of shell grown during the experiment (referred to as shell volume) and the mean thickness of the shell grown during the experiment (referred to as shell thickness) were quantified using micro-CT scanning (Figs 1, 3). The effects of OA on calcification differed across treatments, which was also observed in the transcriptomic response.

**Figure 3.**
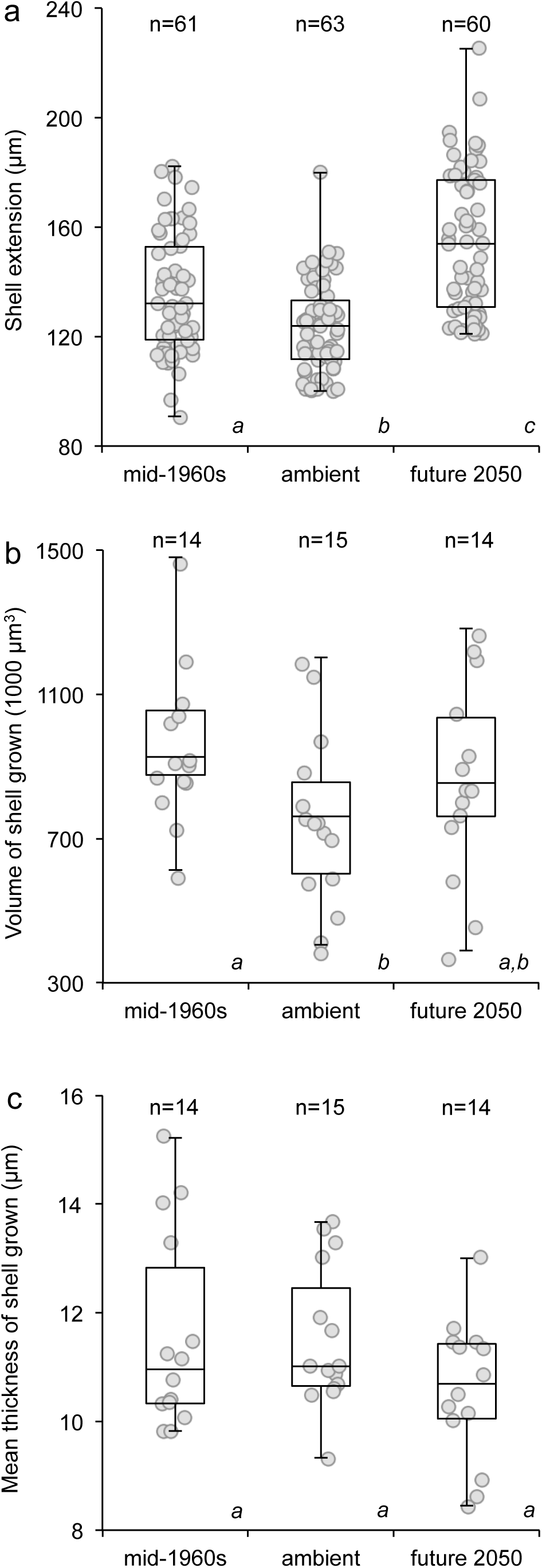
Shell growth of *Atlanta ariejansseni declines* from mid-1960s to ambient ocean pH, but from ambient to future 2050 pH, longer shell was produced. For the boxplots, horizontal lines are median values, boxes are 1^st^ and 3^rd^ quartiles, and bars show the minimum and maximum measurements. Scattered points show all measurements. Significant differences between treatments are denoted by italic letters below each box. (**a**) Shell extension data gathered across pH for 184 specimens (outliers were removed, see methods). (**b**) Shell volume and (**c**) Shell mean thickness grown under different OA scenarios for a randomly selected subset of 43 specimens.

Individuals grew less shell (shorter shell extension and lower volume) under the ambient pH compared to the pH of the mid-1960s, suggesting that ambient conditions in the South Atlantic Ocean already limit calcification of atlantid heteropods. Shell extension and shell volume grown under ambient conditions were found to be significantly lower than shell extension and shell volume grown under the mid-1960s treatment (Fig. 3a, extension Tukey’s HSD p=0.005; volume Mann-Whitney p=0.015). However, shell thickness remained similar between the mid-1960s and ambient treatments (Fig. 3c, Kruskal-Wallis H(2)=2.606, p=0.272). A reduction in calcification from the past to the present has also been found in shelled pteropods from time series data ^38,39^. In the Mediterranean Sea, *Styliola subula* specimens collected in 1921 had thicker shells when compared to specimens collected in 2012, and *Cavolinia inflexa* shells collected in 1910 were denser than those collected in 2012 ^38^. Corresponding reduction in seawater pH in these areas varied by 0.09-0.10 units ^38^. Offshore of northern Australia, a decline in the aragonite saturation state was also found to be accompanied by decreasing shell thickness and increasing shell porosity of two pteropod species from the 1980s to 2009 ^39^. The only perturbation experiment to consider past (higher than ambient) pH on shelled pteropods found no significant differences between shell mass of *Limacina retroversa* grown under pre-industrial (pH 8.2 in that region) and ambient (pH 8.0) conditions ^40^.

A further reduction in calcification with decreasing pH was not observed in *A. ariejansseni* for the 2050 treatment. Instead, a different type of response was found. Individuals grown under future 2050 conditions produced the same volume of shell as individuals grown under ambient conditions (Fig. 3b, volume Mann-Whitney p=0.156), however, shell extension was significantly greater under the future 2050 treatment than under ambient conditions (Fig. 3a, extension Tukey’s HSD p=<0.001). The shells grown under the 2050 treatment were generally thinner (10.6 µm ± 1.3 mean ± s.d.) than those grown under both ambient (11.6 µm ± 1.8) and mid-1960s (11.5 µm ± 1.3) conditions, although this relationship was not significant (Fig. 3c, Mann-Whitney p=0.102). The increased shell extension observed under 2050 conditions in the present study may indicate a stress response to the lowered pH, which is supported by our gene expression results (see below). It has been shown that some calcifying organisms are able to increase the rates of biological processes, such as metabolism and calcification, in response to low pH in order to compensate for the increased acidity ^41^. However, this often comes at a cost to overall fitness ^41^ and such increased rates cannot be sustained in the long term. In pteropods, OA is known to negatively impact metabolic processes ^16,17^.

At the gene expression level, we also found a different response between treatments. From the mid-1960s to the ambient conditions 110 genes were differentially expressed (DE) (66 down- and 44 up-regulated), while from the ambient to the 2050 conditions there were 49 DE genes (12 down and 37 up-regulated), with only 9 of them shared between treatments (Fig. S4, adjusted P < 0.05). In total the DE genes account for approximately 0.5% of the *A. ariejansseni de novo* transcriptome (Table S1), which is in the range of previous transcriptomic responses to high CO_2_ in pteropods (0.001% to 2.6%) ^14,15,49,50^ and copepods (0.25%) ^51^. Based on the transcriptome annotation (Table S2) most genes that were responsive to changes in the carbonate chemistry are potentially involved in the immune response, protein synthesis and degradation, biomineralization, carbohydrate metabolism, morphogenesis and development, ion transport, oxidation-reduction and lipid metabolism (Tables S3-S4).

Most differentially expressed genes identified as potentially involved in biomineralization (Fig. 4) were down-regulated with decreasing pH, along with the observed reduction in overall shell calcification (Fig. 4). Candidate biomineralization genes included those coding for extracellular shell matrix proteins ^52,53^ such as mucins and two chitin binding proteins, but also genes potentially involved in the transport of both proteins and ions to the biomineralization site ^54^, including a sodium dependent transporter and a calcium activated-channel regulator. A smaller fraction of DE biomineralization genes were up-regulated from the mid-1960s to the ambient conditions and/or from the ambient to the future 2050 ocean pH, including members of the mucin, perlucin-like and MAM and LDL-receptor families. These transcriptional changes together with the different calcification responses over changing ocean carbonate chemistry also suggest that distinct genes underlie the control of shell extension and thickness.

**Figure 4.**
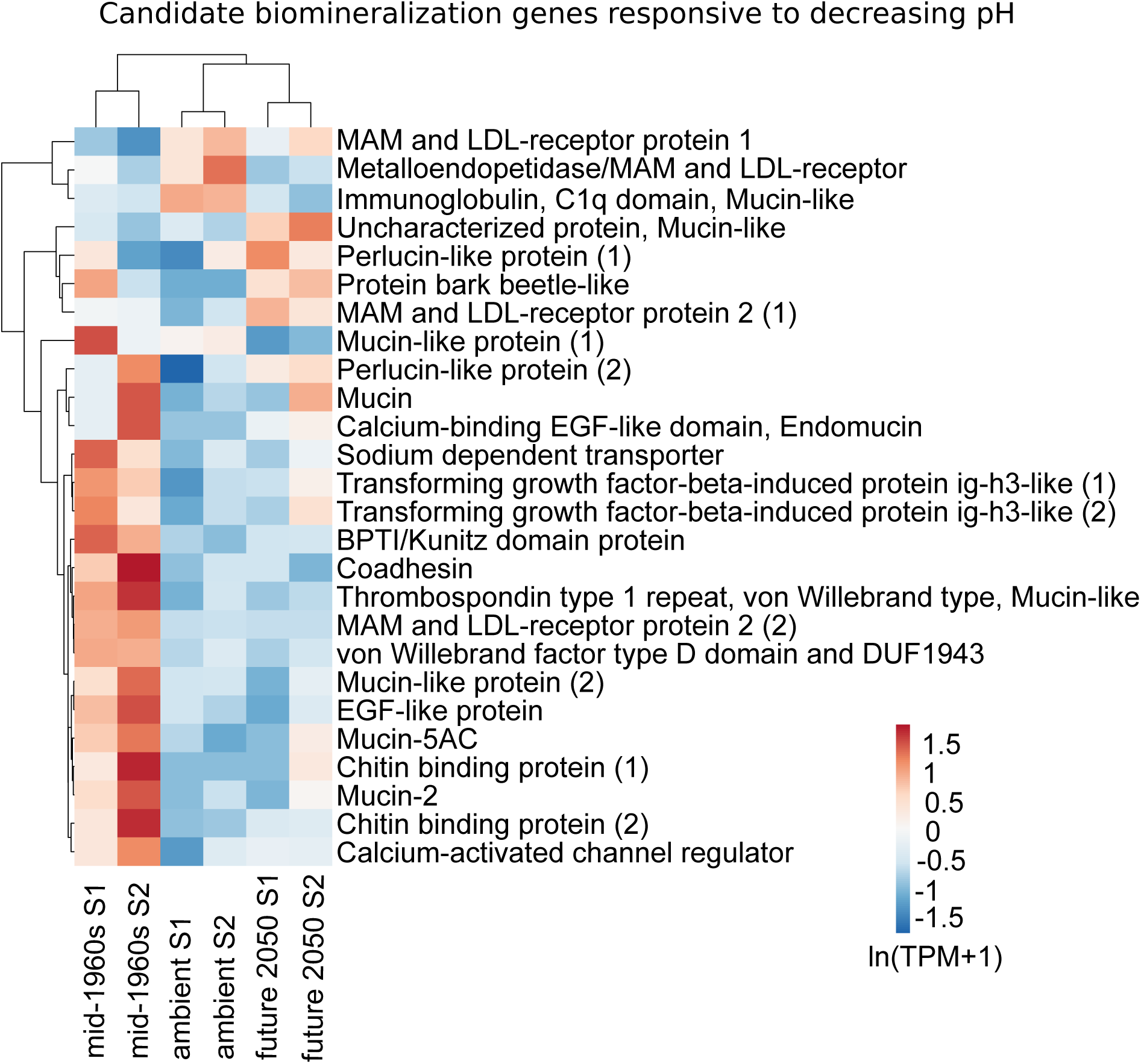
Shell growth and gene expression patterns of candidate biomineralization genes in mid-1960s, ambient and future 2050 ocean pH conditions. Heatmap of candidate biomineralization genes that were responsive to pH changes (adj. p-value < 0.05). Original values of relative abundance of the transcript in units of Transcripts Per Million (TPM) were ln(x + 1)-transformed; pareto scaling was applied to rows. Both rows and columns are clustered using correlation distance and mean linkage.

Differential gene expression analyses indicated a moderate stress response at both ambient and 2050 future system states ^55^ (and references therein), with up-regulation of protein synthesis with decreasing pH (Fig S5a-b in red and Fig. S5d). Such metabolic increase was at the cost of other organism processes with the overall down-regulation of genes involved in development, growth and biomineralization (Fig 4b, Fig. S5a-b in blue and Fig S5c). This response contrasts with the *extreme stress response* ^51,55,56^ observed for the pteropod *Heliconoides inflatus* in a similar 3-day calcification experiment where OA was shown to negatively impact metabolic processes and up-regulate biomineralization ^17^. These differences are likely to be due to taxon-specific responses, although an *extreme stress response* ^55,56^ could have been mitigated by the replete food conditions in our experiment.

### The effects of plentiful food

Plentiful food provided during the incubations may have allowed atlantids to grow thicker shells under experimental conditions in comparison to shell grown *it situ*. OA experiments on shelled pteropods indicate that a lack of food has negative effects on specimen condition ^42^. Nutrition is also likely important for atlantid shell production because they are thought to calcify close to the deep chlorophyll maximum ^27^, a region of higher food availability. Therefore, the juvenile atlantids in OA incubations were all kept under replete food conditions. Specimens were observed to feed well during the experiments, with the chlorophyll-rich algae in their stomachs clearly visible through their transparent shells (Supplementary Figure S1).

Aside from the differences between treatments detailed above, the mean thickness of the shell grown during the OA experiments was significantly higher (1.6 to 2.6 times higher) than the shell grown prior to the experiment for all shells measured in all treatments (t-test, t=-20.720, p=<0.001, n=41). All micro-CT scanned individuals show an initial thickening of the shell (n=43, Fig. 1c,g,i-k) that coincides with the onset of the experiment (apart from two specimens in which the thickening occurred after the onset). It is likely that enhanced calcification from the start of the experiments is related to the availability of plentiful food in all treatments. All of the 3 day OA experimental treatments received the same amount of food, so the effects of food are independent from the effects of the three different treatments in our experiment.

Individuals in the 11 day ambient growth experiments were fed a much higher concentration of food because it was anticipated that they would remain in the experiment for up to 24 days (the experiment was terminated at 11 days due to cannibalism caused by metamorphosis). When comparing these specimens at day three (n=8, pH 8.15) to specimens incubated for three days under the same ambient conditions for the OA experiment (n=63, pH 8.14 ± 0.02), but with eight times less food, a significant difference in the shell extension was found (t-test, t=8.714, p=<0.001). Specimens given a higher concentration of food grew on average 1.62 times longer shell than those with the lower food concentration (mean 208 µm and 129 µm respectively).

These results suggest that increased food availability leads to increased calcification, and that a plentiful food supply could offset OA induced reduction in calcification. Similar trends have been found in other calcifying organisms, including pteropods^43^, corals ^44,45^ and the early benthic stages of the bivalve mollusc *Mytilus edulis* ^46^. The pteropod *Heliconoides inflatus* was found to produce shells that were 40% thicker and 20% larger in diameter during periods of naturally high nutrient concentrations in the Cariaco basin (compared to specimens sampled during oligotrophic conditions) ^43^. A review of OA studies on calcifying marine organisms found that an intermediate or high food supply increased the resistance to low pH for growth and calcification ^47^.

### A complex response to OA

The results of this first study on the effects of OA upon atlantid heteropods have revealed a complex organismal response. Ocean carbonate chemistry with decreasing pH had a negative, but varied effect on calcification, both from the higher pH of the mid-1960s to the ambient conditions, and from the ambient conditions to lower ocean pH predicted for 2050. A reduction in shell extension and shell volume from the mid-1960s to the present conditions suggests that ambient water chemistry is already limiting atlantid calcification. An increase in shell extension from ambient to 2050 conditions may indicate a stress response to grow to a larger size as quickly as possible with rapidly decreasing pH. Gene expression analyses indicated a moderate stress response at both ambient and 2050 future conditions ^48^, with down-regulation of growth and calcification, and up-regulation of protein synthesis with decreasing pH. However, the high availability of food may have increased calcification across all treatments, and it may be that increased food supply can mitigate some of the negative effects of OA on juvenile atlantids. At the adult stage, however, atlantids feed primarily on shelled pteropods ^22^, and the assumed decline in the abundance of OA sensitive pteropods will have a negative effect on food availability for adult atlantids. The fairly short time that it takes an atlantid to reach maturity may mean that multiple generations are produced each year, and this could help atlantids adapt more quickly to a rapidly changing ocean ^49^.

In summary, the findings of this study indicate that calcification in atlantids is already impacted by OA in the South Atlantic Ocean, however, some of the effects of OA on atlantid calcification could be mitigated by replete food. Our results demonstrate that the effects of OA on atlantid calcification and their subsequent export of carbonate to deep waters is not straight forward, and likely depends on whether these organisms are able to survive and maintain calcification under stressful conditions in the long term. Evidence suggests that both shelled pteropods and atlantids survived the Cretaceous-Paleogene extinction event (KPg or KT) and Paleocene-Eocene Thermal Maximum (PETM), both periods of extreme perturbation in the ocean’s carbon cycle ^50,51^. This observation gives some hope that aragonite shelled holoplanktonic gastropods will be able to adapt to our changing oceans, even though the rate of change is unprecedented relative to the geological record. *Atlanta ariejansseni* resides in cool convergence waters where rapid changes in water temperature and water stratification are expected to be additional stressors ^52^. Future studies should seek to understand the synergistic effects of ocean acidification and warming, to understand variability in the environment in which atlantids live (and environmental tolerances that they may already have), and to thoroughly investigate how nutrition affects calcification and growth.

## Methods

### Specimen collection and staining

Specimens of *Atlanta ariejansseni* were collected in the Southern Subtropical Convergence Zone during the Atlantic Meridional Transect (AMT) 27 (DY084/085) cruise of the *RRS Discovery*. Animals for growth experiments were collected on the 24^th^ October 2017 at 35°58 S, 27°57 W and for OA experiments on the 26^th^ October 2017 at 41°09 S, 30°00 W. For both experiments, samples were collected using a 1 m diameter ring net with 200 µm mesh and a closed cod-end for three slow, short (20 minute) oblique tows to a maximum depth of 100 m. Samples were collected during hours of darkness between 00:38 and 01:57. Specimens of *A. ariejansseni* were immediately sorted from the net samples using a light microscope and placed in calcein indicator for two hours in the dark (MERCK Calcein indicator for metal determination, CAS 1461-15-0, concentration 50 mg/l in seawater filtered through a 0.2 µm filter). Specimens were then gently rinsed with 0.2 µm filtered seawater and introduced into the experimental carboys.

### OA experiment

Surface seawater was filtered at 0.2 µm into four 60 litre barrels, which underwent the following treatments. In one barrel, lowered pH was achieved by bubbling 795 ppm CO_2_ in air through the water for 12 hours, attaining 0.11 pH units below ambient (pH 8.05). In a second barrel, higher pH was achieved by bubbling 180 ppm CO_2_ in air through the water for 12 hours, attaining 0.02 pH units above ambient (pH 8.18). The final two barrels of ambient and control (ambient) water at pH 8.16 ± 0.00 were not subjected to any gas bubbling. During gas bubbling, all water was maintained at ambient ocean temperatures (at the depth of collection), between 14-16 °C within a temperature controlled room on board the *RRS Discovery*. Temperature, salinity and pH (resolution of pH 0.001, precision of pH ± 0.002, HANNA HI5522-02) were measured from the four barrels after 12 hours, and samples to measure Dissolved Inorganic Carbon (DIC) concentration were collected. Immediately prior to specimen collection, three carboys of six litres were filled for each of the treated and ambient seawaters. A further two carboys were filled with ambient seawater to act as controls (no specimens added). Natural algal concentrations at the deep chlorophyll maximum in the study region are ∼0.2 µg/l (AMT data extracted from British Oceanographic Data Centre). To ensure that food was not limiting calcification rates, dried algae (a mixture of 33.3% *Phaeodactylum*, 33.3% *Nannochloropsis*, 33.3% *Tetraselmis*) was added to each of the carboys (including the controls) at a concentration of 0.6 mg/l (0.2 mg/l/day; 3.6 mg per carboy).

Calcein-stained specimens (n=274, 217 for physical measurements, 57 for gene expression analysis) were introduced into the carboys in a random order. Between 90 and 92 juveniles were exposed to each treatment. Specimens of *A. ariejansseni* were identified by their shell morphology ^31^, and juveniles were recognised based on size, presence of the velum and absence of black eye pigmentation. Carboys were sealed and immediately incubated at ambient temperature within a temperature controlled room (14-16 °C). Blackout fabric was draped over the carboys to maintain low light levels. The carboys were incubated for three days. At the end of the third day, temperature, salinity and pH were measured, and DIC concentration samples were collected from all carboys. Specimens were removed from the carboys, and examined under a light microscope to verify that they were still alive (movement). 20 live juveniles were pooled for each treatment (from one replicate), preserved in RNAlater (Invitrogen) and frozen for RNAseq analyses. The remaining specimens were flash frozen in liquid nitrogen and stored at -20 °C until analysis.

Across all treatments, the pH of the experimental carboys was stable from the start to the end of the experiment and remained fairly consistent between replicates (Table 2). The pH of two control carboys containing ambient water and no specimens also remained stable over the three days and did not differ from the ambient experiment. The water remained supersaturated with regards to aragonite throughout all treatments (Table 2) and no signs of shell dissolution were observed (surface etching or clouding of the shells). Mortality was extremely low across all treatments, with only a single specimen (ambient treatment) having died during the experiment.

**Table 2.** Measured and calculated (using CO2SYS) carbonate system parameters for the three ocean acidification treatments and the resulting shell growth of *Atlanta ariejansseni* (averaged across all replicates). N1 is the number of specimens measured for the shell extension (total n=184) and N2 is the number of specimens measured for volume, thickness and diameter (n=43). Values are presented ± 1 s.d. when averaged over replicates.

| Sample | Day | Mean measured DIC (μmol/kg) | Mean calculated TA (μmol/kg) | Mean measured pH (NBS) | Mean calculated pCO<sub>2</sub> | Mean calculated ΩAr. | N1 | Mean shell extension (μm) | N2 | Mean volume of shell grown (1000 μm<sup>3</sup>) | Mean thickness of shell pre-exp (μm) | Mean thickness of shell grown (μm) | Mean maximum shell diameter (μm) |
| --- | --- | --- | --- | --- | --- | --- | --- | --- | --- | --- | --- | --- | --- |
| Ambient control | 0 | 2112 | 2338 | 8.16 | 410 | 2.51 | - | - | - | - | - | - | - |
| Ambient control | 3 | 2103 ± 2 | 2308 ± 0 | 8.13 ± 0.00 | 441 ± 9 | 2.27 ± 0.02 | - | - | - | - | - | - | - |
| Mid-1960s | 0 | 2110 | 2352 | 8.18 | 391 | 2.65 | - | - | - | - | - | - | - |
| Mid-1960s | 3 | 2109 ± 6 | 2341 ± 5 | 8.19 ± 0.02 | 378 ± 18 | 2.53 ± 0.10 | 61 | 136 ± 22 | 14 | 963 ± 210 | 5.81 ± 0.89 | 11.58 ± 1.81 | 400 ± 49 |
| Ambient | 0 | 2110 | 2339 | 8.16 | 405 | 2.53 | - | - | - | - | - | - | - |
| Ambient | 3 | 2139 ± 10 | 2350 ± 2 | 8.14 ± 0.02 | 425 ± 20 | 2.33 ± 0.08 | 63 | 124 ± 16 | 15 | 755 ± 246 | 5.96 ± 0.61 | 11.51 ± 1.30 | 361 ± 72 |
| Future 2050 | 0 | 2150 | 2329 | 8.05 | 553 | 2.05 | - | - | - | - | - | - | - |
| Future 2050 | 3 | 2154 ± 13 | 2312 ± 11 | 8.03 ± 0.00 | 564 ± 5 | 1.84 ± 0.02 | 60 | 155 ± 26 | 14 | 871 ± 270 | 5.68 ± 0.73 | 10.58 ± 1.29 | 386 ± 96 |

### Growth rate experiment

Surface seawater was filtered at 0.2 µm and maintained at ambient temperature in a 6 l carboy. Temperature, salinity and pH were measured and samples for DIC concentration were collected prior to adding specimens. Dried algae were added to the carboy at a concentration of 4.8 mg/l, to allow for approximately 24 days of feeding (0.2 mg/l/day. 28.8 mg per carboy). Calcein-stained specimens (n=57) were introduced into the carboy, which was immediately sealed and incubated at ambient temperature, covered with blackout fabric. After three days, temperature, salinity and pH measurements were made, and a sample for DIC concentration was collected. Up to ten live individuals were removed and flash frozen in liquid nitrogen and stored at -20 °C until analysis. Any dead specimens retrieved at this stage were removed from the experiment and discarded (total for the whole experiment n=13). Subsequent to sampling, the carboy was sealed and returned to ambient, dark conditions. Sampling was carried out in the same way for water parameters and specimens approximately every two days. The experiment was terminated at 11 days because some specimens metamorphosed and began to cannibalise other animals in the experiment.

### Water chemistry

Dissolved Inorganic Carbon (DIC) samples were filtered into 5 ml glass vials. The water samples contained no head space and were poisoned with 15 µl of saturated mercury (II) chloride (HgCl_2_). Analysis of DIC was carried out at the Royal Netherlands Institute for Sea Research (NIOZ), Texel, The Netherlands, using a Technicon Traacs 800 autoanalyzer spectrophotometric system following the methodology of Stoll et al. ^53^. pH was measured on the NBS scale using a research grade benchtop pH meter (HANNA HI5522-02) and a glass electrode with a resolution of 0.001 units, and a precision of ± 0.002 units. The pH meter was regularly calibrated using NBS standards. Other carbonate system parameters were calculated from the measured DIC and measured pH ^54^ using CO2SYS (Excel V2.3) ^55^. The calculation used the constants K1 and K2 from Mehrbach et al. ^56^ refitted by Dickson and Millero ^57^ and the KHSO4 dissociation constant of Dickson ^58^. Nutrient concentrations (P and Si) were measured from surface CTD samples in the regions where water was collected for the experiments ^59^. All carbonate system parameters are correlated (Pearson r= -0.996–0.977, p=<0.002), except DIC and total alkalinity (Pearson r= -0.114, p=0.687).

### Shell extension

Shell cleaning and fluorescent imaging was carried out at the Royal Netherlands Institute for Sea Research (NIOZ), Texel, The Netherlands. Organic material was removed from the shells by oxidising the specimens in a Tracerlab low temperature (∼100 °C) asher. This ensured minimal damage compared to chemical and physical washing techniques. Specimens were air dried for 24 hours and then oxidised in the low temperature asher for five hours. Specimens were then gently rinsed with ethanol and ultra-high purity water (MilliQ) to remove any ash residue, and dried in a cool oven (40 °C) for 15 minutes. Specimens were imaged using a Zeiss Axioplan 2 microscope with a Colibri light source and filter (excitation 485/20, FT 510, emission 515-565) producing a final wavelength of 515-565 nm. The extent of fluorescent shell was measured along the suture between the last whorl and the preceding whorl using the software FIJI (ImageJ) ^60^. Despite extreme care being taken during shell handling, the growing edge of some shells was damaged, providing only a minimum measure of shell extension. Therefore, for the OA experiments, severely damaged specimens (n=31), where the shell edge was broken back to the calcein stained region (start of the experiment), were not included in subsequent analyses.

### Shell thickness and volume

Specimens were visually inspected using light microscopy to determine whether there was any damage at the growing edge of the shell. Between 14 and 15 undamaged specimens were randomly selected from each treatment (total n=43 specimens) and scanned using a Zeiss Xradia 520 Versa microCT at Naturalis Biodiversity Center, Leiden, The Netherlands. Scans were carried out using between 140/10 and 150/10 kV/W for between 2 and 3 hours per specimen. The scan resolution was 0.54–0.68 µm with an exposure time of 5–10 seconds. Data were processed using the software Avizo 2019.1 (Thermo Fisher). Shells were segmented to separate the part of the shell that had grown during the experiment. This was achieved by manually matching the fluorescent images to the microCT thickness map using whorl counting and other landmarks such as growth lines and repair marks on the shells (Fig. 1a-h). The segmented shells were then analysed for the volume and mean thickness of the shell grown during the experiment. MicroCT images were also measured to determine maximum shell diameter. Mean shell thickness was found to negatively correlate to shell diameter (Pearson r=-0.669, p=<0.001, n=39), but only for shell grown prior to the experiments (Fig. S2). The mean thickness of shell grown during the experiments was not correlated to shell diameter for any of the treatments.

### Statistical analyses

To determine correlations between variables, for example between shell thickness and shell diameter, Pearson’s Chi-squared was used, and to confirm correlations between the carbonate system parameters, a full pairwise matrix of Pearson’s correlation coefficients was made. Growth measurement data for the OA experiments (shell extension, shell thickness and shell volume), and shell extension per day for the ambient growth experiment were checked for a normal distribution using a Levene’s test. Shell extension data were normally distributed (Levene’s p=<0.001). The shell thickness and shell volume were not normally distributed (Levene’s p=0.253 and p=0.663, respectively). To identify whether shell extension from the OA experiment varied between treatments, a one-way ANOVA was performed, followed by a Tukey’s HSD posthoc test to indicate more detailed differences between treatments. A Kruskal-Wallace test for equal medians was carried out on the shell thickness and shell volume from the OA experiments, and on the shell extension from the ambient growth experiment. Where a significant difference between sample medians was identified, a Mann-Whitney posthoc test was performed to indicate more detailed differences between treatments. Statistical analyses were carried out using PAST v3.12 ^61^.

### RNA extraction and sequencing

Due to the small size of *Atlanta ariejansseni* juveniles (mean diameter of a subset 381 ± 73 µm, n=39), total RNA was extracted from samples of 8–10 individuals randomly pooled from each treatment using the RNeasy Plus Micro Kit (QIAGEN). This sampling provided 2 replicates per treatment (6 samples in total). Each sample of total RNA was analyzed for quantity and quality using the Bioanalyzer 2100 (Agilent Technologies) with a RNA 6000 Nano Chip. All samples had RIN scores ranging from 7.0 to 9.4 and were used for library preparation and sequencing. Libraries (n=6) were generated with the NEBNext® Ultra II Directional Library Prep Kit for Illumina (New England BioLabs) using the manufacturer’s protocol for Poly(A) mRNA magnetic isolation from 1 µg total RNA per sample. Total RNA was added to NEBNext Sample Purification beads to isolate the mRNA. Purified mRNA was then fragmented into approx. 300 base-pair (bp) fragments and reverse transcribed into cDNA using dUTPs in the synthesis of the second strand. cDNA fragments were size selected and amplified with 8-9 PCR cycles using NEBNEXT Multiplex Dual Index kit (New England BioLabs) according to the manufacturer’s instructions. Libraries were checked for quantity and quality on Bioanalyzer 2100 using an Agilent DNA High Sensitivity Chip. Average library sizes of 420 up to 450 bps (∼300 bp insert +128 bp sequencing adapters) were accessed using the Agilent Bioanalyzer 2100. Sequencing was performed at the BaseClear BV Leiden on an Illumina NovaSeq 6000 platform using paired-end 150 base-pair sequences. All six libraries were sequenced producing a minimum of 6 giga base pairs (Gb) per library.

### *De novo* assembly and data analysis

Raw reads were processed using trimmomatic (version 0.38 ^62^) to remove adapter sequences and reads lower than 36 bps, and checked for quality using FastQC (version 0.11.8). Trimmed reads were pooled and assembled with Trinity v2.8.4 ^63^ using default parameters. Open reading frames (ORFs) of the *de novo* transcriptome assembly were predicted using Transdecoder v5.5.0 ^63^. The ORFs of the longest isoforms from each trinity locus were blasted against a subset of the NCBI nr database (release from 9/20/19) including all Mollusca (txid6447), Stramenophiles (txid33634) and Viridiplantae (txid33090). Contigs having a best hit with molluscan sequences (e-value <10e^-5^) were considered *bona fide Atlanta ariejansseni* transcripts; all the other contigs (for example derived from the mixture of algae fed to the animals or other potential contaminants) were removed from the assembly. After this filtering, the distribution of GC content in the assembly appeared unimodal (Fig. S3) suggesting the major sources of contamination were removed without compromising the transcriptome completeness as determined with BUSCO ^64^ (Table S1). Next, transcript quantifications were estimated based on the raw reads and the raw transcriptome assembly as reference, using Salmon ^65^. Only quantifications of transcripts present in the clean transcriptome assembly were used in differential gene expression estimation using the DESeq2 package ^66^ in pairwise comparisons between the ambient *vs*. higher mid-1960s pH and lower 2050 *vs*. ambient pH. Significant differentially expressed genes were selected based on P-adj values < 0.05 corrected for multiple testing with the Benjamini-Hochberg procedure, which controls false discovery rate (FDR) (Tables S3 and S4). Annotation of the clean transcriptome assembly was performed using the Trinotate v3.2.0 pipeline, which levered the results from different functional annotation strategies including homology searches using BLAST+ against Swissprot (release October 2019) and protein domain detection using HMMER ^67^ against PFAM ^68^ (release September 2018). Gene ontology (GO) terms obtained from this annotation strategy (Table S2) were trimmed using the GOSlimmer tool ^69^ followed by enrichment analyses using the GO-MWU method described in ^70^. This method used adaptive clustering of GO categories and Mann–Whitney U tests ^71^ based on ranking of signed log p-values to identify over‐ represented GO terms in the categories “Biological Process” and “Molecular Function”. In addition, genes were grouped in 10 main categories (*i.e*. putative processes or other) according to their BLAST+ best hits in RefSeq or Swissprot (releases October 2019) and associated GO terms (Table S3, Table S4): ‘immune response’, ‘protein synthesis’, ‘protein degradation’, ‘biomineralization’, ‘carbohydrate metabolism’, ‘development/morphogenesis’, ‘ion transport’, ‘oxidation-reduction’, ‘lipid metabolism’ and ‘other’. Gene expression heatmaps with hierarchical clustering of expression profiles were created with ClustVis ^72^.

## Data availability

All data supporting the findings of this study is provided in the online supplementary information. Raw reads used in this study were deposited at NCBI BioProject PRJNA17165. The Transcriptome Shotgun Assembly has been deposited at DDBJ/EMBL/GenBank under the accession GIOD00000000. The version described in this paper is the first version, GIOD01000000.

## Acknowledgements

We are very grateful to Rob Langelaan and Dirk van der Marel (Naturalis) for microCT scanning specimens, to Vassilis Kitidis (Plymouth Marine Laboratory) and Matthew Humphreys (NIOZ) for discussion on carbonate chemistry and checking carbonate system calculations, and to the captain, crew and scientists who took part in cruise DY084/085 (AMT27) onboard the *RRS Discovery* (PSO: A. Rees). Plankton collection on the AMT27 cruise was funded by a Vidi grant (016.161351) from the Netherlands Organisation for Scientific Research (NWO) to KTCAP. The Atlantic Meridional Transect is funded by the UK Natural Environment Research Council through its National Capability Long-term Single Centre Science Programme, Climate Linked Atlantic Sector Science (grant number NE/R015953/1). This study contributes to the international IMBeR project and is contribution number 335 of the AMT programme. LKD was supported by the Netherlands Earth System Science Centre (NESSC), Grant Number: 024.002.001 from the Dutch Ministry of Education, Culture and Science. This project has received funding from the European Union’s Horizon 2020 research and innovation programme under the Marie Sklodowska-Curie grant agreement No 746186 [POSEIDoN, DW-P] and grant agreement No 844345 [EPIC, PRS].

## Author contributions

DW-P, LM, KTCAP and PRS designed the study, DW-P, LM, KTCAP and EG performed the research, DW-P, LD, KB, PRS and ED carried out sample preparation and analysis. DW-P, LM and PRS carried out data analysis. All authors contributed to manuscript preparation.

## Supplementary Tables and Figures provided separately

Table S1. Transcriptome assembly statistics of *Atlanta ariejansseni* reported using Trinity v2.8.4, Quast (Galaxy version 4.6.3), and BUSCO v3 before and after filtering for potential contaminant contigs. The *Atlanta ariejansseni de novo* transcriptome assembly consisted of 28,512 predicted ‘genes’ (here genes as defined by ^73^). Giving overall good quality and 93.5% completeness, the *A. ariejansseni* transcriptome is suitable for differential expression analysis, and comparable to prior pteropod *de novo* transcriptomes ^16,17^.

Table S2. Transcriptome functional annotation of *Atlanta ariejansseni* using Trinotate: summary of the strategies and statistics.

Table S3. Genes responsive to the high pH treatment grouped by potential functional categories. Differential gene expression was performed using the DESeq2 based in the pairwise comparison present vs. past (i.e. ambient vs. high pH).

Table S4. Genes responsive to the low pH treatment grouped by potential functional categories. Differential gene expression was performed using the DESeq2 package based in the pairwise comparison future vs. present (i.e. low pH vs. ambient).

Figure S1. (**a**) Fluorescence image showing a repair (indicated by white arrow) to the side of a shell that is fluorescing (mid-1960s treatment, replicate 3) and (**b**) a cross section of the same specimen imaged using microCT showing the repair from the inside of the shell. (**c, d**) Bright green algae were visible in the stomachs of the specimens, for example specimens from the ‘normal’ rate of growth experiment, day 9 (**c**), and specimens from the mid-1960s treatment, replicate 2 (**d**).

Figure S2. The relationship between maximum shell diameter and shell thickness of *Atlanta ariejansseni*. (**a**) The thickness of shell grown prior to the experiment shows a significant negative correlation to the maximum shell diameter, indicating that shell becomes thinner as the specimen increases in size. (**b**) The thickness of shell grown during the OA experiments is not related to maximum shell diameter because the normal growth was altered by varying pH and an increase in food concentration. (**c**)Mean thickness of shell grown prior to the experiment significantly correlates to the mean thickness of the shell grown during the experiment. (Pearson r=0.687, p=<0.001). So, the specimens that grew thicker shells before the experiments grew proportionally thicker shells during the experiments. This demonstrates that individuals also naturally vary in their ability to calcify, and these individual differences persist across the changes in environmental conditions that they experienced within our experiments.

Figure S3. Distribution of GC content before and after filtering contaminant sequences.

Figure S4. Venn diagram representing the overlap of genes differentially expressed in *Atlanta ariejansseni* juveniles in the different pH treatments; (≥1.5-fold change; Benjamini-Hochberg-adjusted P<0.05).

Figure S5. Overview of the gene expression response of *Atlanta ariejansseni* juveniles to high and low ocean pH. (**a, b**) Hierarchical clustering of gene ontology terms enriched by genes up-regulated (red) or down-regulated (blue) and summarized by molecular function (MF) and biological process (BP) for each pairwise comparison: ambient *vs* mid-1960s, and future 2050 *vs* ambient. GO categories associated with protein synthesis were consistently up-regulated with decreasing pH: translation (GO:0006412), structural constituents of the ribosome (GO:0003735), RNA binding (GO:0003723), ribosome biogenesis (GO:0042254), and ribonucleoprotein complex assembly (GO:0022618). On the other hand, GO categories associated with morphogenesis and organismal development were down-regulated under decreasing pH: locomotion (GO:0040011), cell morphogenesis (GO:0000902), cell adhesion (GO:0007155), nervous system process (GO: GO:0050877) and cell differentiation (GO: GO:0030154). The size of the font indicates the significance of the term as indicated by the inset key. The fraction preceding the GO term indicates the number of genes annotated with the term that pass an unadjusted p-value threshold of 0.05. Heatmap of the (**c**) fraction of genes involved in morphogenesis and development and (**d**) fraction of genes involved in protein synthesis that were responsive to pH changes (adj. p-value < 0.05). Original values of relative abundance of the transcript in units of Transcripts Per Million (TPM) were ln(x+1)-transformed; pareto scaling was applied to rows. Both rows and columns are clustered using correlation distance and mean linkage.

## References

1. Bednaršek, N., Možina, J., Vogt, M., O’Brien, C. & Tarling, G. A. The global distribution of pteropods and their contribution to carbonate and carbon biomass in the modern ocean. Earth System Science Data 4, 167–186 (2012).

2. Manno, C. et al. Threatened species drive the strength of the carbonate pump in the northern Scotia Sea. Nature Communications 9, (2018).

3. Buitenhuis, E. T., Le Quéré, C., Bednaršek, N. & Schiebel, R. Large Contribution of Pteropods to Shallow CaCO 3 Export. Global Biogeochemical Cycles 33, 458–468 (2019).

4. Bathmann, U. V., Noji, T. T. & von Bodungen, B. Sedimentation of pteropods in the Norwegian Sea in autumn. Deep Sea Research Part A. Oceanographic Research Papers 38, 1341–1360 (1991).

5. Berner, R. A. & Honjo, S. Pelagic sedimentation of aragonite: its geochemical significance. Science 211, 940–942 (1981).

6. Gruber, N. et al. The oceanic sink for anthropogenic CO 2 from 1994 to 2007. Science 363, 1193–1199 (2019).

7. Pachauri, R. K. et al. Climate change 2014: synthesis report. Contribution of Working Groups I, II and III to the fifth assessment report of the Intergovernmental Panel on Climate Change. (R. Pachauri and L. Meyer (editors), Geneva, Switzerland, IPCC, 2014).

8. Riebesell, U. & Gattuso, J.-P. Lessons learned from ocean acidification research. Nature Climate Change 5, 12 (2014).

9. Gattuso, J.-P. et al. Contrasting futures for ocean and society from different anthropogenic CO 2 emissions scenarios. Science 349, aac4722 (2015).

10. Zeebe, R. E., Ridgwell, A. & Zachos, J. C. Anthropogenic carbon release rate unprecedented during the past 66 million years. Nature Geoscience 9, 325 (2016).

11. Kitidis, V., Brown, I., Hardman-Mountford, N. & Lefèvre, N. Surface ocean carbon dioxide during the Atlantic Meridional Transect (1995–2013); evidence of ocean acidification. Progress in Oceanography 158, 65–75 (2017).

12. Bednaršek, N., Harvey, C. J., Kaplan, I. C., Feely, R. A. & Možina, J. Pteropods on the edge: Cumulative effects of ocean acidification, warming, and deoxygenation. Progress in Oceanography 145, 1–24 (2016).

13. Kroeker, K. J. et al. Impacts of ocean acidification on marine organisms: quantifying sensitivities and interaction with warming. Global Change Biology 19, 1884–1896 (2013).

14. Bednaršek, N. et al. Extensive dissolution of live pteropods in the Southern Ocean. Nature Geoscience 5, 881–885 (2012).

15. Manno, C. et al. Shelled pteropods in peril: Assessing vulnerability in a high CO 2 ocean. Earth-Science Reviews 169, 132–145 (2017).

16. Maas, A. E., Lawson, G. L., Bergan, A. J. & Tarrant, A. M. Exposure to CO 2 influences metabolism, calcification and gene expression of the thecosome pteropod *Limacina retroversa*. The Journal of Experimental Biology 221, jeb164400 (2018).

17. Moya, A. et al. Near-future pH conditions severely impact calcification, metabolism and the nervous system in the pteropod *Heliconoides inflatus*. Global Change Biology 22, 3888–3900 (2016).

18. Bednaršek, N. et al. New ocean, new needs: Application of pteropod shell dissolution as a biological indicator for marine resource management. Ecological Indicators 76, 240–244 (2017).

19. Burridge, A. K. et al. Diversity and abundance of pteropods and heteropods along a latitudinal gradient across the Atlantic Ocean. Progress in Oceanography 158, 213–223 (2017).

20. Batten, R. L. & Dumont, M. P. Shell ultrastructure of the Atlantidae (Heteropoda, Mesogastropoda) *Oxygyrus* and *Protatlanta*, with comments on *Atlanta inclinata*. Bulletin of the American Museum of Natural History 157, 263–310 (1976).

21. Fabry, V. J. Shell growth rates of pteropod and heteropod molluscs and aragonite production in the open ocean: implications for the marine carbonate system. Journal of Marine Research 48, 209–222 (1990).

22. Lalli, C. M. & Gilmer, R. W. Pelagic snails: the biology of holoplanktonic mollusks. (Stanford University Press, 1989).

23. Mucci, A. The solubility of calcite and aragonite in seawater at various salinities, temperatures, and one atmosphere total pressure. American Journal of Science 283, 780–799 (1983).

24. Gazeau, F. et al. Impact of elevated CO 2 on shellfish calcification. Geophysical Research Letters 34, (2007).

25. Comeau, S., Gorsky, G., Alliouane, S. & Gattuso, J.-P. Larvae of the pteropod *Cavolinia inflexa* exposed to aragonite undersaturation are viable but shell-less. Marine Biology 157, 2341–2345 (2010).

26. Caldeira, K. & Wickett, M. E. Anthropogenic carbon and ocean pH. Nature 425, 365–365 (2003).

27. Wall-Palmer, D. et al. Vertical distribution and diurnal migration of atlantid heteropods. Marine Ecology Progress Series 587, 1–15 (2018).

28. Negrete-García, G., Lovenduski, N. S., Hauri, C., Krumhardt, K. M. & Lauvset, S. K. Sudden emergence of a shallow aragonite saturation horizon in the Southern Ocean. Nature Climate Change 9, 313–317 (2019).

29. Fabry, V. J., McClintock, J. B., Mathias, J. T. & Grebmeier, J. M. Ocean acidification at high latitudes: the bellwether. Oceanography 22, 160–171 (2009).

30. Wall-Palmer, D. et al. A review of the ecology, palaeontology and distribution of atlantid heteropods (Caenogastropoda: Pterotracheoidea: Atlantidae). Journal of Molluscan Studies 82, 221–234 (2016).

31. Wall-Palmer, D., Burridge, A. K. & Peijnenburg, K. T. C. A. *Atlanta ariejansseni,* a new species of shelled heteropod from the Southern Subtropical Convergence Zone (Gastropoda, Pterotracheoidea). ZooKeys 604, 13–30 (2016).

32. Lischka, S., Büdenbender, J., Boxhammer, T. & Riebesell, U. Impact of ocean acidification and elevated temperatures on early juveniles of the polar shelled pteropod *Limacina helicina*: mortality, shell degradation, and shell growth. Biogeosciences 8, 919–932 (2011).

33. Peck, V. L., Oakes, R. L., Harper, E. M., Manno, C. & Tarling, G. A. Pteropods counter mechanical damage and dissolution through extensive shell repair. Nature Communications 9, (2018).

34. Bednaršek, N., Tarling, G. A., Fielding, S. & Bakker, D. C. E. Population dynamics and biogeochemical significance of *Limacina helicina antarctica* in the Scotia Sea (Southern Ocean). Deep Sea Research Part II: Topical Studies in Oceanography 59–60, 105–116 (2012).

35. Hunt, B. P. V. et al. Pteropods in Southern Ocean ecosystems. Progress in Oceanography 78, 193–221 (2008).

36. Roberts, D. et al. Interannual pteropod variability in sediment traps deployed above and below the aragonite saturation horizon in the Sub-Antarctic Southern Ocean. Polar Biology 34, 1739–1750 (2011).

37. Hartin, C. A., Bond-Lamberty, B., Patel, P. & Mundra, A. Ocean acidification over the next three centuries using a simple global climate carbon-cycle model: projections and sensitivities. Biogeosciences 13, 4329–4342 (2016).

38. Howes, E. L., Eagle, R. A., Gattuso, J.-P. & Bijma, J. Comparison of Mediterranean Pteropod Shell Biometrics and Ultrastructure from Historical (1910 and 1921) and Present Day (2012) Samples Provides Baseline for Monitoring Effects of Global Change. PLOS ONE 12, e0167891 (2017).

39. Roger, L. M. et al. Comparison of the shell structure of two tropical Thecosomata (*Creseis acicula* and *Diacavolinia longirostris*) from 1963 to 2009: potential implications of declining aragonite saturation. ICES Journal of Marine Science 69, 465–474 (2012).

40. Manno, C., Morata, N. & Primicerio, R. *Limacina retroversa’s* response to combined effects of ocean acidification and sea water freshening. Estuarine, Coastal and Shelf Science 113, 163–171 (2012).

41. Wood, H. L., Spicer, J. I. & Widdicombe, S. Ocean acidification may increase calcification rates, but at a cost. Proceedings of the Royal Society B: Biological Sciences 275, 1767–1773 (2008).

42. Howes, E. L. et al. Sink and swim: a status review of thecosome pteropod culture techniques. Journal of Plankton Research 36, 299–315 (2014).

43. Oakes, R. L. & Sessa, J. A. Determining how biotic and abiotic variables affect the shell condition and parameters of *Heliconoides inflatus* pteropods from a sediment trap in the Cariaco Basin. Biogeosciences 17, 1975–1990 (2020).

44. Cohen, A. L. & Holcomb, M. Why corals care about ocean acidification: uncovering the mechanism. Oceanography 22, 118–127 (2009).

45. Holcomb, M., McCorkle, D. C. & Cohen, A. L. Long-term effects of nutrient and CO2 enrichment on the temperate coral *Astrangia poculata* (Ellis and Solander, 1786). Journal of Experimental Marine Biology and Ecology 386, 27–33 (2010).

46. Thomsen, J., Casties, I., Pansch, C., Körtzinger, A. & Melzner, F. Food availability outweighs ocean acidification effects in juvenile *Mytilus edulis* : laboratory and field experiments. Global Change Biology 19, 1017–1027 (2013).

47. Ramajo, L. et al. Food supply confers calcifiers resistance to ocean acidification. Scientific Reports 6, (2016).

48. Sokolova, I. M., Frederich, M., Bagwe, R., Lannig, G. & Sukhotin, A. A. Energy homeostasis as an integrative tool for assessing limits of environmental stress tolerance in aquatic invertebrates. Marine Environmental Research 79, 1–15 (2012).

49. Peijnenburg, K. T. C. A. & Goetze, E. High evolutionary potential of marine zooplankton. Ecology and Evolution 3, 2765–2781 (2013).

50. Peijnenburg, K. T. C. A. et al. The origin and diversification of pteropods predate past perturbations in the Earth’s carbon cycle. bioRxiv (2019) doi: 10.1101/813386.

51. Wall-Palmer, D. et al. Fossil-calibrated molecular phylogeny of atlantid heteropods (Gastropoda, Pterotracheoidea). BMC Evolutionary Biology (in review).

52. Swart, N. C., Gille, S. T., Fyfe, J. C. & Gillett, N. P. Recent Southern Ocean warming and freshening driven by greenhouse gas emissions and ozone depletion. Nature Geoscience 11, 836–841 (2018).

53. Stoll, M. H. C., Bakker, K., Nobbe, G. H. & Haese, R. R. Continuous-Flow Analysis of Dissolved Inorganic Carbon Content in Seawater. Analytical Chemistry 73, 4111–4116 (2001).

54. Hoppe, C. J. M., Langer, G., Rokitta, S. D., Wolf-Gladrow, D. A. & Rost, B. Implications of observed inconsistencies in carbonate chemistry measurements for ocean acidification studies. Biogeosciences 9, 2401–2405 (2012).

55. Pierrot, D. E., Wallace, D. W. R. & Lewis, E. MS Excel Program Developed for CO2 System Calculations. (2011) doi: 10.3334/CDIAC/otg.CO2SYS_XLS_CDIAC105a.

56. Mehrbach, C., Culberson, C. H., Hawley, J. E. & Pytkowicx, R. M. Measurement of the apparent dissociation constants of carbonic acid in seawater at atmospheric pressure. Limnology and Oceanography 18, 897–907 (1973).

57. Dickson, A. G. & Millero, F. J. A comparison of the equilibrium constants for the dissociation of carbonic acid in seawater media. Deep Sea Research Part A. Oceanographic Research Papers 34, 1733–1743 (1987).

58. Dickson, A. G. Thermodynamics of the dissociation of boric acid in synthetic seawater from 273.15 to 318.15 K. Deep Sea Research Part A. Oceanographic Research Papers 37, 755–766 (1990).

59. Woodward, E. M. S. & Harris, C. Atlantic Meridional Transect cruise AMT27 (DY084) micro-molar nutrient measurements from CTD bottle samples during 2017. (2019) doi: 10.5285/915141EE-46C9-4DA1-E053-6C86ABC09800.

60. Schindelin, J. et al. Fiji: an open-source platform for biological-image analysis. Nature Methods 9, 676–682 (2012).

61. Hammer, Ø., Harper, D. A. T. & Ryan, P. D. PAST: paleontological statistics software package for education and data analysis. Palaeontologia Electronica 4, 1–9 (2001).

62. Bolger, A. M., Lohse, M. & Usadel, B. Trimmomatic: A flexible trimmer for Illumina sequence data. Bioinformatics 30, 2114–2120 (2014).

63. Haas, B. J. et al. De novo transcript sequence reconstruction from RNA-seq using the Trinity platform for reference generation and analysis. Nature Protocols 8, 1494–1512 (2013).

64. Simão, F. A., Waterhouse, R. M., Ioannidis, P., Kriventseva, E. V. & Zdobnov, E. M. BUSCO: Assessing genome assembly and annotation completeness with single-copy orthologs. Bioinformatics 31, 3210–3212 (2015).

65. Patro, R., Duggal, G., Love, M. I., Irizarry, R. A. & Kingsford, C. Salmon provides fast and bias-aware quantification of transcript expression. Nature methods 14, 417–419 (2017).

66. Love, M. I., Huber, W. & Anders, S. Moderated estimation of fold change and dispersion for RNA-seq data with DESeq2. Genome biology 15, 550 (2014).

67. Eddy, S. R. Accelerated profile HMM searches. PLoS Computational Biology 7, (2011).

68. El-Gebali, S. et al. The Pfam protein families database in 2019. Nucleic Acids Research 47, D427–D432 (2019).

69. Faria, D. GOSlimmer. (2017).

70. Wright, R. M., Aglyamova, G. V., Meyer, E. & Matz, M. V. Gene expression associated with white syndromes in a reef building coral, Acropora hyacinthus. BMC Genomics 16, 371 (2015).

71. Voolstra, C. R. et al. Rapid Evolution of Coral Proteins Responsible for Interaction with the Environment. PLoS ONE 6, e20392 (2011).

72. Metsalu, T. & Vilo, J. ClustVis: a web tool for visualizing clustering of multivariate data using Principal Component Analysis and heatmap. Nucleic Acids Research 43, W566–W570 (2015).

73. Haas, B. J. et al. De novo transcript sequence reconstruction from RNA-seq using the Trinity platform for reference generation and analysis. Nature Protocols 8, 1494 (2013).

