## Supplementary Tables and Figures for "Current and future ocean chemistry negatively impacts calcification in predatory planktonic snails"

Table S1. Transcriptome assembly statistics of *Atlanta ariejansseni* reported using Trinity v2.8.4, Quast (Galaxy version 4.6.3), and BUSCO v3 before and after filtering for potential contaminant contigs. The *Atlanta ariejansseni de novo* transcriptome assembly consisted of 28,512 predicted 'genes' (here genes as defined by [73]). Giving overall good quality and 93.5% completeness, the *A. ariejansseni* transcriptome is suitable for differential expression analysis, and comparable to prior pteropod *de novo* transcriptomes [16,17].

| | **Raw assembly** | **Filtered assembly** |
|---|---|---|
| **Trinity** | | |
| # Predicted genes (locus) | 352,021 | 28,512 |
| # Transcripts (contigs) | 766,227 | 97,483 |
| Total size (bp) | 400,030,081 | 109,082,979 |
| **Quast** | | |
| Largest contig (bp) | 18,325 | 18,325 |
| GC (%) | 43,34 | 49,65 |
| N50 (bp) | 1442 | 2001 |
| **BUSCO (%)** | | |
| Complete | **94.4** | **93.5** |
| Complete and single-copy | 91.2 | 90.3 |
| Complete and duplicated | 3.2 | 3.2 |
| Fragmented | 3.5 | 3.8 |
| Missing | 2.1 | 2.7 |

Table S2. Transcriptome functional annotation of *Atlanta ariejansseni* using Trinotate: summary of the strategies and statistics.

| Annotation strategy | Annotated genes |
|---|---|
| BLASTX/Swissprot (E-value < 1.00E-3) | 21,494 (75%) |
| BLASTP/Swissprot (E-value < 1.00E-3) | 22,183 (78%) |
| HMMER/PFAM (E-value < 1.00E-5) | 19,339 (68%) |
| Gene Ontology assignments from best blast hit (E -value< 1.00E-50) and PFAM | 14,828 (52%) |

Table S3. Genes responsive to the high pH treatment grouped by potential functional categories. Differential gene expression was performed using the DESeq2 based in the pairwise comparison present vs. past (i.e. ambient vs. high pH).

Provided separately as an Excel file

Table S4. Genes responsive to the low pH treatment grouped by potential functional categories. Differential gene expression was performed using the DESeq2 package based in the pairwise comparison future vs. present (i.e. low pH vs. ambient).

Provided separately as an Excel file

### Supplementary Figures

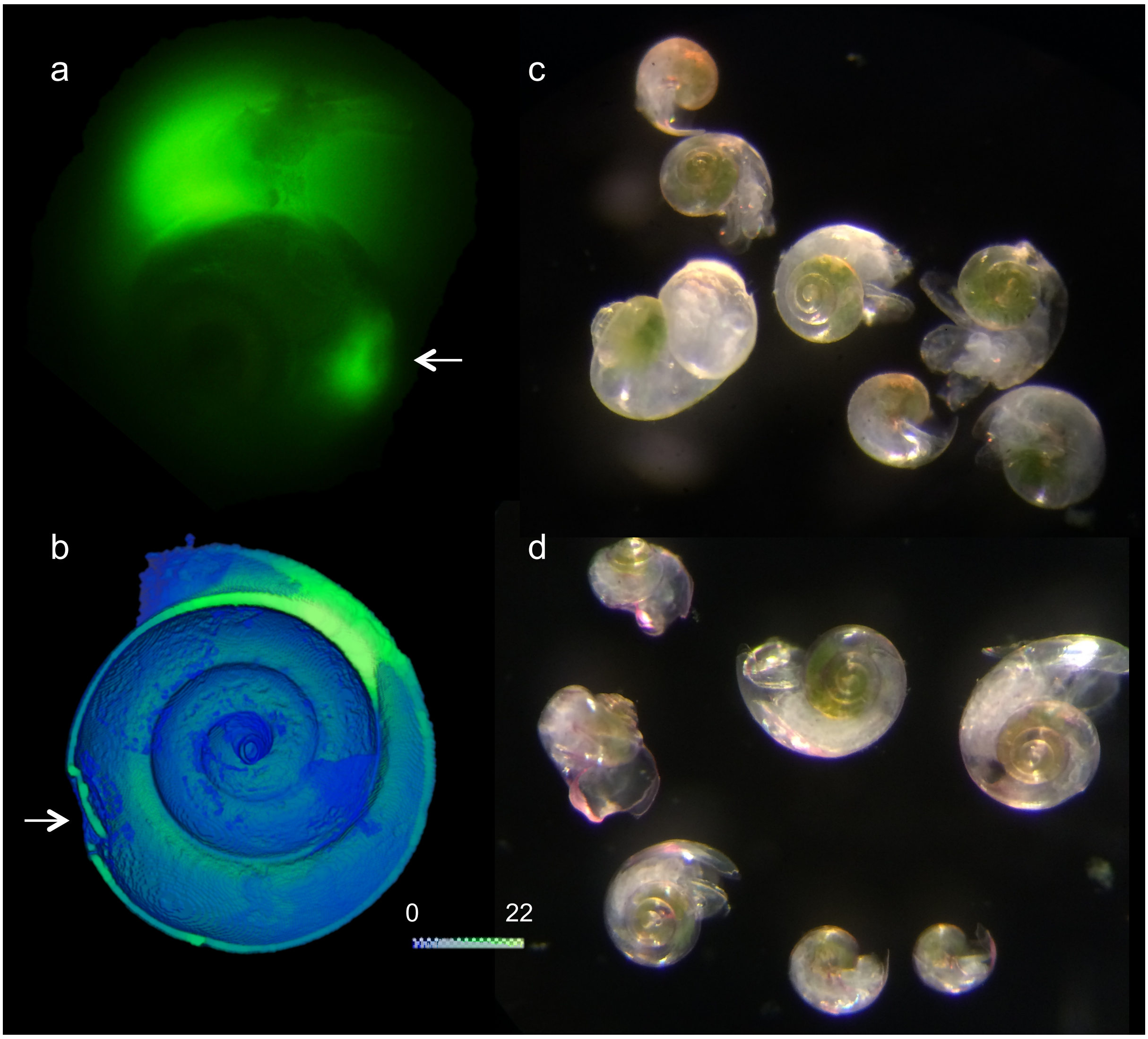


Figure S1. (**a**) Fluorescence image showing a repair (indicated by white arrow) to the side of a shell that is fluorescing (mid-1960s treatment, replicate 3) and (**b**) a cross section of the same specimen imaged using microCT showing the repair from the inside of the shell. (**c, d**) Bright green algae were visible in the stomachs of the specimens, for example specimens from the 'normal' rate of growth experiment, day 9 (**c**), and specimens from the mid-1960s treatment, replicate 2 (**d**).

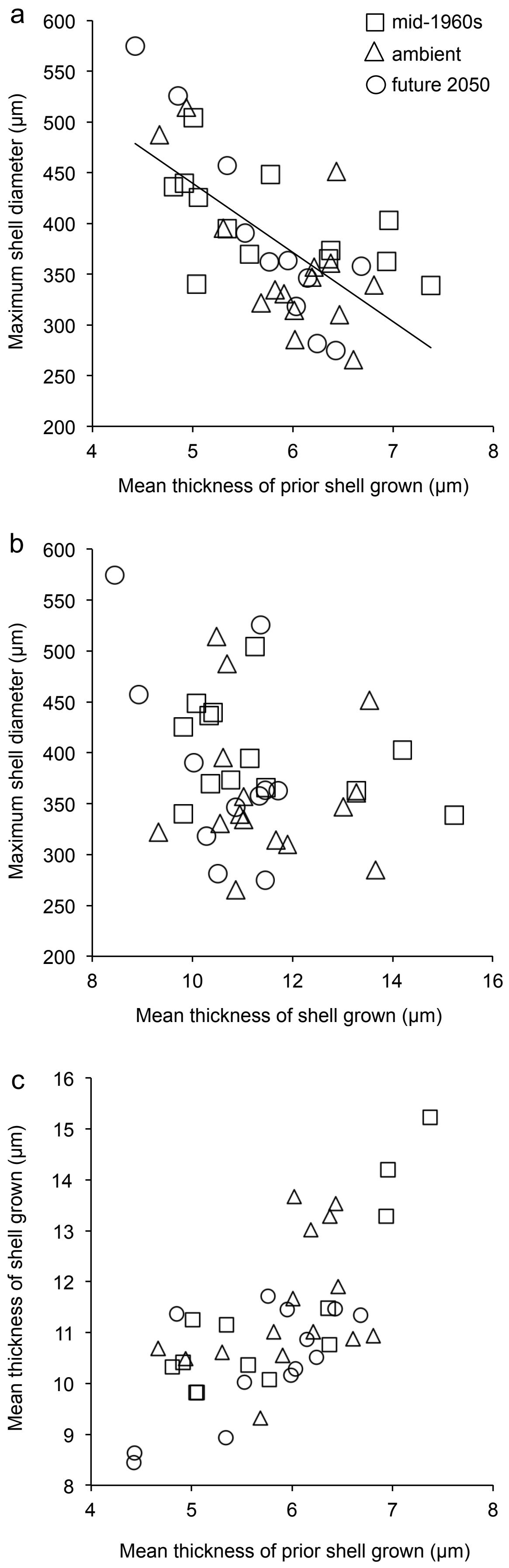
a
mid-1960s
ambient
future 2050
Maximum shell diameter (μm)
Mean thickness of prior shell grown (μm)
b
Maximum shell diameter (μm)
Mean thickness of shell grown (μm)
c
Mean thickness of shell grown (μm)
Mean thickness of prior shell grown (μm)

Figure S2. The relationship between maximum shell diameter and shell thickness of *Atlanta ariejansseni*. (**a**) The thickness of shell grown prior to the experiment shows a significant negative correlation to the maximum shell diameter, indicating that shell becomes thinner as the specimen increases in size. (**b**) The thickness of shell grown during the OA experiments is not related to maximum shell diameter because the normal growth was altered by varying pH and an increase in food concentration. (**c**) Mean thickness of shell grown prior to the experiment significantly correlates to the mean thickness of the shell grown during the experiment. (Pearson r=0.687, p=<0.001). So, the specimens that grew thicker shells before the experiments grew proportionally thicker shells during the experiments. This demonstrates that individuals also naturally vary in their ability to calcify, and these individual differences persist across the changes in environmental conditions that they experienced within our experiments.

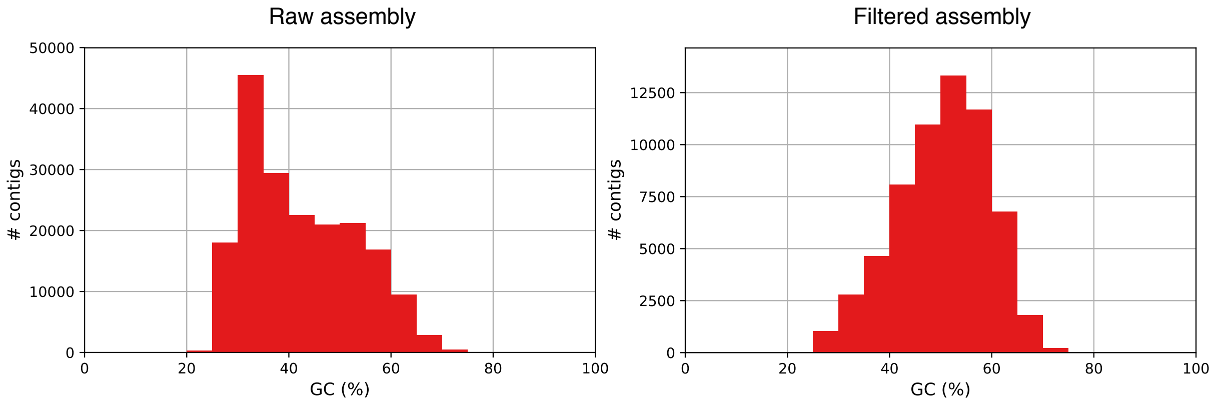


Figure S3. Distribution of GC content before and after filtering contaminant sequences.

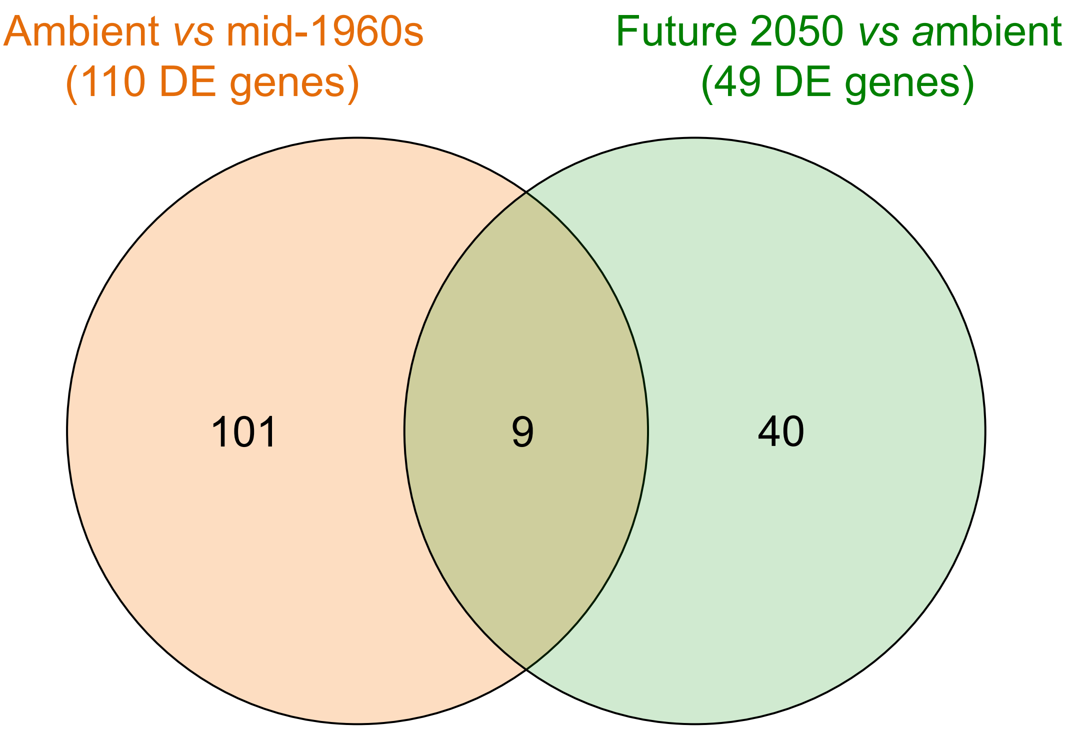


Figure S4. Venn diagram representing the overlap of genes differentially expressed in *Atlanta ariejansseni* juveniles in the different pH treatments; (≥1.5-fold change; Benjamini-Hochberg-adjusted $P<0.05$).

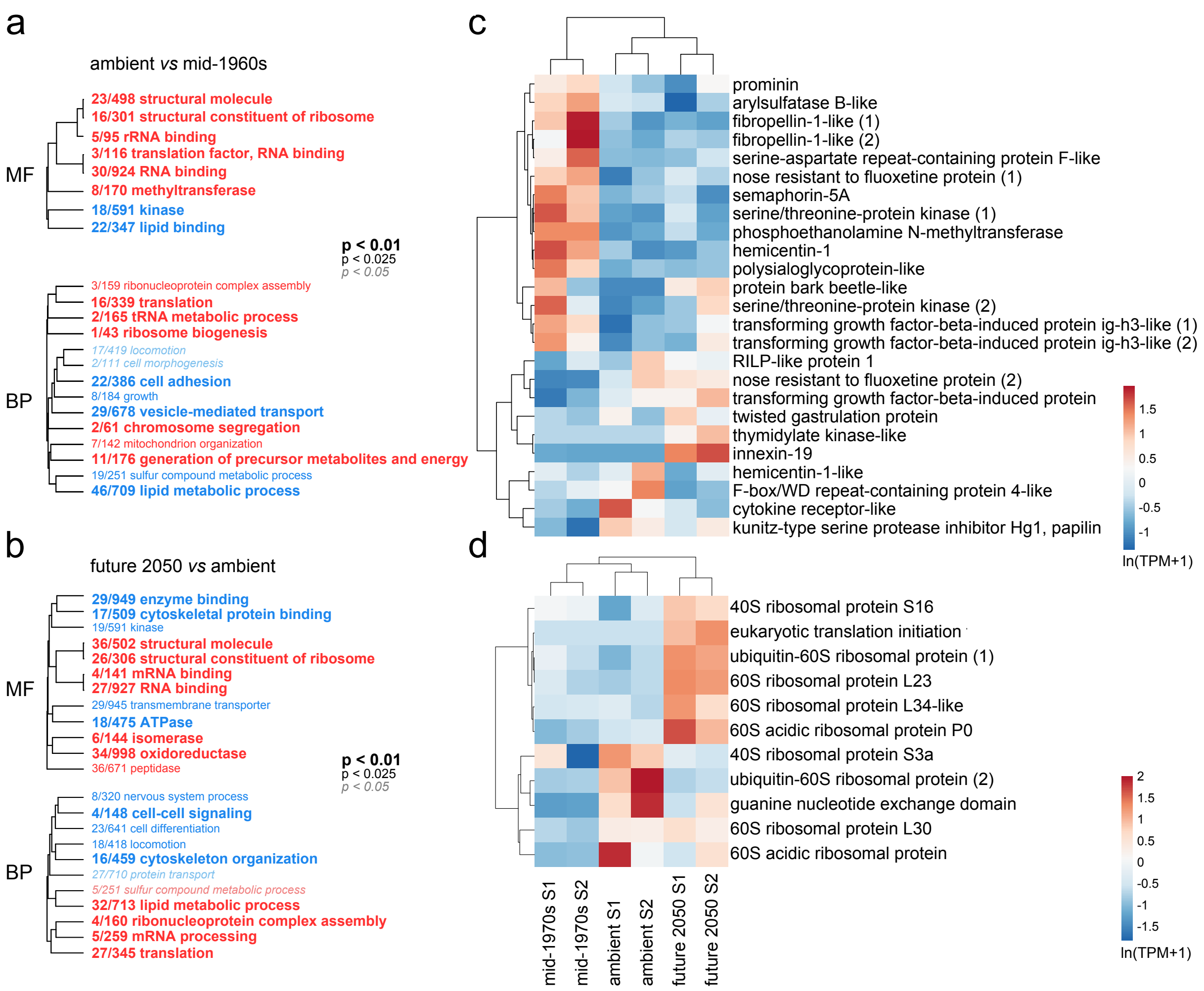


Figure S5. Overview of the gene expression response of *Atlanta ariejansseni* juveniles to high and low ocean pH. (**a, b**) Hierarchical clustering of gene ontology terms enriched by genes up-regulated (red) or down-regulated (blue) and summarized by molecular function (MF) and biological process (BP) for each pairwise comparison: ambient *vs* mid-1960s, and future 2050 *vs* ambient. GO categories associated with protein synthesis were consistently up-regulated with decreasing pH: translation (GO:0006412), structural constituents of the ribosome (GO:0003735), RNA binding (GO:0003723), ribosome biogenesis (GO:0042254), and ribonucleoprotein complex assembly (GO:0022618). On the other hand, GO categories associated with morphogenesis and organismal development were down-regulated under decreasing pH: locomotion (GO:0040011), cell morphogenesis (GO:0000902), cell adhesion (GO:0007155), nervous system process (GO: GO:0050877) and cell differentiation (GO: GO:0030154). The size of the font indicates the significance of the term as indicated by the inset key. The fraction preceding the GO term indicates the number of genes annotated with the term that pass an unadjusted p-value threshold of 0.05. Heatmap of the (**c**) fraction of genes involved in morphogenesis and development and (**d**) fraction of genes involved in protein synthesis that were responsive to pH changes (adj. p-value < 0.05). Original values of relative abundance of the transcript in units of Transcripts Per Million (TPM) were ln(x+1)-transformed; pareto scaling was applied to rows. Both rows and columns are clustered using correlation distance and mean linkage.
